# Coordinated Cross-Talk Between the Myc and Mlx Networks in Liver Regeneration and Neoplasia

**DOI:** 10.1101/2021.08.05.455215

**Authors:** Huabo Wang, Jie Lu, Frances Alencastro, Alexander Roberts, Julia Fiedor, Patrick Carroll, Robert N. Eisenman, Sarangarajan Ranganathan, Michael Torbenson, Andrew W. Duncan, Edward V. Prochownik

## Abstract

**Background & Aims:** The c-Myc (Myc) bHLH-ZIP transcription factor is deregulated in most cancers. In association with Max, Myc controls target genes that supervise metabolism, ribosome biogenesis, translation and proliferation. This “Myc Network” cross-talks with the “Mlx Network”, which consists of the Myc-like proteins MondoA and ChREBP and Max-like Mlx. Together, this “Extended Myc Network” regulates both common and distinct genes targets. Here we studied the consequence of *Myc* and/or *Mlx* ablation in the liver, particularly those pertaining to hepatocyte proliferation, metabolism and spontaneous tumorigenesis.

**Methods:** We examined the ability of hepatocytes lacking *Mlx* (*Mlx*KO) or *Myc+Mlx* (double KO or DKO) to repopulate the liver over an extended period of time in a murine model of Type I tyrosinemia. We also compared this and other relevant behaviors, phenotypes and transcriptomes of the livers to those from previously characterized *Myc*KO, *Chrebp*KO and *Myc*KO x *Chrebp*KO mice.

**Results:** Hepatocyte regenerative potential deteriorated as the Extended Myc Network was progressively dismantled. Genes and pathways dysregulated in *Mlx*KO and DKO hepatocytes included those pertaining to translation, mitochondrial function and non-alcoholic fatty liver disease (NAFLD). The Myc and Mlx Networks were shown to cross-talk, with the latter playing a disproportionate role in target gene regulation. All cohorts also developed NAFLD and molecular evidence of early steatohepatitis. Finally, *Mlx*KO and DKO mice displayed extensive hepatic adenomatosis.

**Conclusions:** In addition to demonstrating cooperation between the Myc and Mlx Networks, this study revealed the latter to be more important in maintaining proliferative, metabolic and translational homeostasis, while concurrently serving as a suppressor of benign tumorigenesis.

**Synopsis:** The Myc and Mlx Networks exhibit extensive cross-talk and regulate distinct but overlapping sets of transcriptional targets. The current work demonstrates the cooperation between these two Networks in supporting the regenerative capabilities of normal hepatocytes while also revealing that the Mlx Network serves as a suppressor of spontaneous hepatic adenomatosis

## Introduction

c-Myc (Myc) is a bHLH-Zip transcription factor that regulates numerous target genes, which collectively support survival, proliferation, metabolism, ribosome biogenesis and translation [1-8]. Positive regulation involves Myc’s direct sequence-specific DNA binding in heterodimeric association with its obligate bHLH-Zip partner, Max [6, 7]. This occurs at canonical “E-box” elements that typically reside in the vicinity of proximal promoters [9-12]. Bound Myc-Max heterodimers recruit an assortment of transcription co-factors and chromatin modifiers such as histone acetylases and methyltransferases that collectively increase chromatin accessibility, relieve transcriptional pausing and increase the rate and efficiency of mRNA elongation [13-17]. Down-regulation of these genes, often occurring during cellular quiescence or differentiation, involves a reduction in Myc levels and a shift to E-box occupancy by heterodimers now comprised of Max and members of the transcriptionally repressive bHLH-ZIP “Mxd family” that includes Mxd1-4 and the less related Mnt and Mga factors [3, 18-20]. Together, their binding reverses the chromatin modifications mediated by Myc-Max binding and restores transcriptional repression. Negative regulation by Myc is more indirect and involves interaction with and inhibition of positively-acting transcription factors such as Miz1 and Sp1 [21, 22]. The loss of transcriptional balance maintained by these different competing interactions is a feature of transformed cells, which often deregulate and/or over-express Myc [4, 10, 14, 23].

The “Myc Network” cross-talks and shares considerable regulatory overlap with a structurally related but distinct group of bHLH-Zip transcription factors that comprise the so-called “Mlx Network” [1-3, 8, 24, 25][20]. Classically believed to control target gene sets smaller and more functionally restricted than those overseen by Myc, the Myc-like equivalents of the Mlx Network include the transcription factors ChREBP and MondoA. Upon binding glucose and other nutrients, these cytoplasmic proteins translocate to the nucleus, heterodimerize with the Max-like protein Mlx and bind to their target genes at “carbohydrate response elements” (ChoREs) comprised of tandem E-boxes separated by 5-6 nucleotides [26, 27]. Myc Network and Mlx Network members (collectively termed the “Extended Myc Network”) can bind to one another’s DNA target sequences and some genes are dually regulated by both sets of factors; however the numbers of these genes and the degree to which their regultation is a result of binding to shared versus separate sites has not been clearly delineated [28-32]. Although the Mlx Network is less widely implicated in tumorigenesis than the Myc Network, recurrent *MLX* gene deletions nevertheless occur in as many as 10-20% of several human cancers, with the precise fraction correlating with the size of the deletion (https://portal.gdc.cancer.gov/genes/ENSG00000108788) [1-3, 13, 33][20].

We previously explored the roles for these two networks in normal hepatocyte proliferation utilizing mice lacking the enzyme fumarylacetoacetate hydrolase (FAH). These animals serve as a model for Type I hereditary tyrosinemia in which FAH’s absence allows toxic tyrosine catabolites to accumulate, leading to hepatic necrosis and liver failure [34-36]. Treatment with the drug 2-[2-nitro-4-trifluoromethylbenzoyl]-1,3-cyclohexanedione (NTBC) blocks the enzyme 4-hydroxyphenylpyruvic dioxygenase, which catalyzes the second step in tyrosine catabolism, thus preventing the accumulation of these deleterious intermediates and circumventing the lethal consequences of FAH deficiency. Immuno-compromised FRG-NOD (*Fah*^−/−^) mice can thus be used as a robust and sensitive animal model in which to evaluate the regenerative potential of any other hepatocyte population, so long as it is *Fah+/+*. Cells are delivered intrasplenically followed by the cyclic withdrawal and reinstatement of NTBC over several months. As recipient hepatocytes accumulate toxic tyrosine intermediates and die, they are replaced by the donor cells, which expand as much as 50-100-fold before eventually comprising up to 70% of the hepatic mass and allowing the recipients to achieve NTBC-independence [25, 37]. The FAH model thus places greater proliferative demands on regenerating hepatocytes than does two-thirds partial hepatectomy (PH), which represents the gold standard for liver regeneration [38]. It also permits the simultaneous delivery of two or more competing populations of hepatocytes to the same recipient, thus allowing for a direct comparison of their relative proliferative rates within the identical environment.

Using this approach, we previously showed that wild-type (WT) and *Myc-/-* (*Myc*KO) hepatocytes possess indistinguishable regenerative potential [37]. This is quite different from most other cases where Myc’s loss in either non-transformed or transformed cells or tissues profoundly suppresses proliferation [25, 37, 39-43]. In contrast, the proliferation of *Chrebp-/-* (*Chrebp*KO) hepatocytes has been show to be significantly impaired and *Myc*KO x *Chrebp*KO hepatocytes are even more defective [25]. These findings indicated that normal hepatocyte regeneration is more dependent upon the Mlx Network than the Myc Network and that the two pathways cross-talk and rescue one another’s defects to varying degrees. At the same time, they raise questions about the possible functional redundancy of MondoA in the context of ChREBP’s loss.

We have now further explored the relationship between the Myc and Mlx Networks by generating two additional mouse strains. In the first (hereafter referred to as “*Mlx*KO”), deletion of the *Mlx* gene functionally inactivates the entire Mlx Network, including any potential rescue by MondoA that might have existed in *Chrebp*KO mice [25]. The second mouse strain contains a “double knockout” of both *Myc* and *Mlx* (hereafter “DKO”) that further inactivates the Extended Myc Network. We show that hepatocytes from both strains, but particularly the latter, are profoundly compromised in repopulating the livers of *Fah-/-* recipients. They also show markedly attenuated expression of genes that are direct targets for both the Myc and Mlx Networks and that control mitochondrial structure and function, ribosomal biogenesis and more general aspects of mRNA processing and translation. Older mice of both groups also develop nonalcoholic fatty liver disease (NAFLD) akin to that previously described in *Myc*KO, *Chrebp*KO or *Myc*KO x *Chrebp*KO mice. Finally and unexpectedly, over one-third of older *Mlx*KO and DKO mice develop multi-focal hepatic adenomas occasionally associated with small regions of hepatocellular carcinoma (HCC). These results further support the idea that the Myc and Mlx Networks cross-talk and cooperatively regulate a range of pathways related to energy metabolism, lipid balance, translation and proliferation. Finally, they reveal a heretofore unsuspected role for the Mlx Network as a suppressor of benign hepatic adenomatosis [44].

## Results

### Repopulation by MlxKO and DKO hepatocytes is severely compromised

Donor mouse strains employed for competitive hepatocyte repopulation studies carried homozygous “floxed” alleles of the *Mlx* and/or *Myc* genes (Fig. 1A & B and Table 1) and expressed an albumin promoter-driven Tamoxifen-inducible CreER [25, 45]. Five daily i.p. injections of Tamoxifen were sufficient to allow inactivation of each allele by the time hepatocytes were transplanted 3-4 months later (Fig. 1C).

**Table 1.**
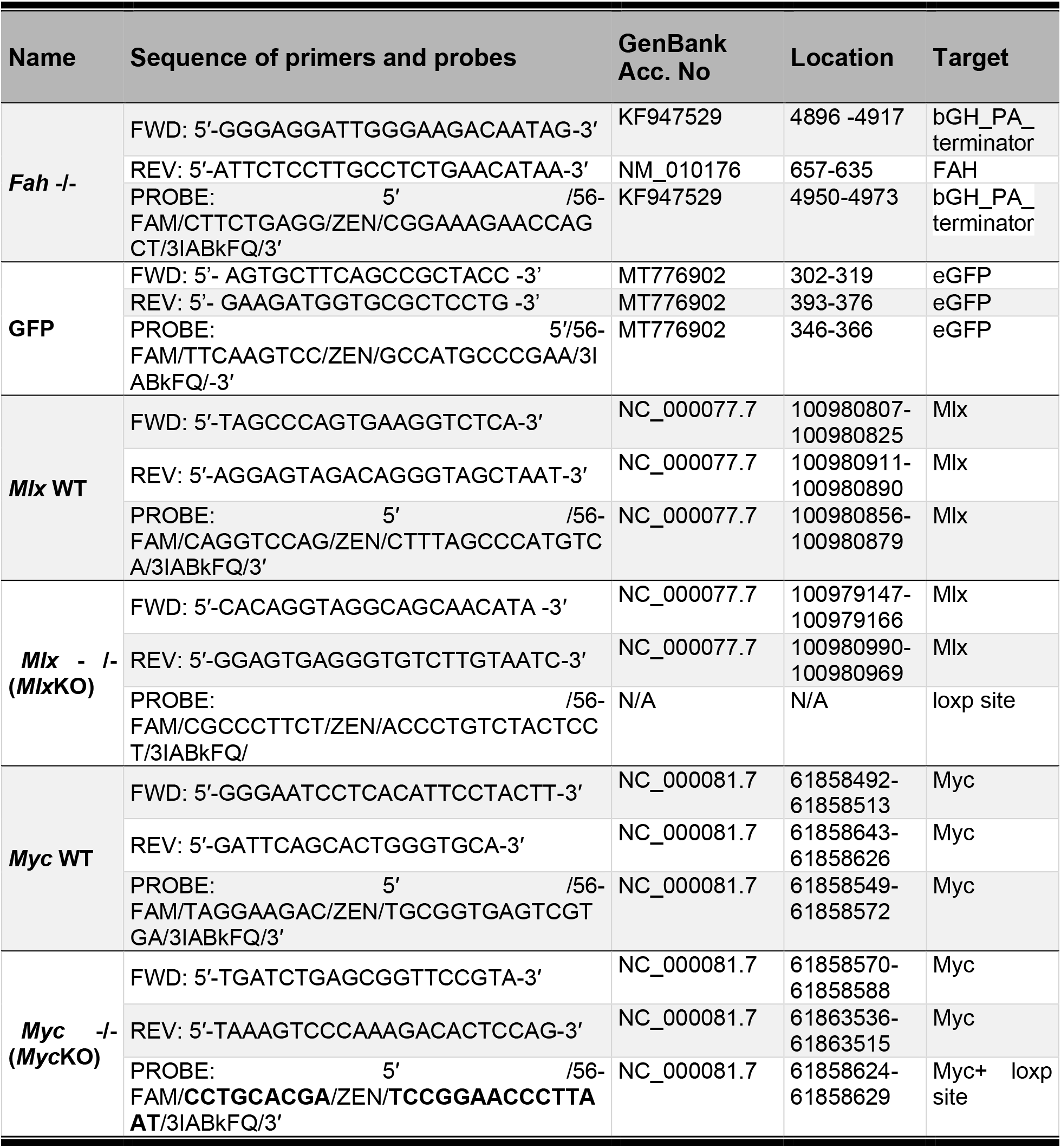
qPCR primers and probes used to quantify each of alleles shown in Fig. 1A and B and other necessary genes as indicated.

**Fig. 1.**
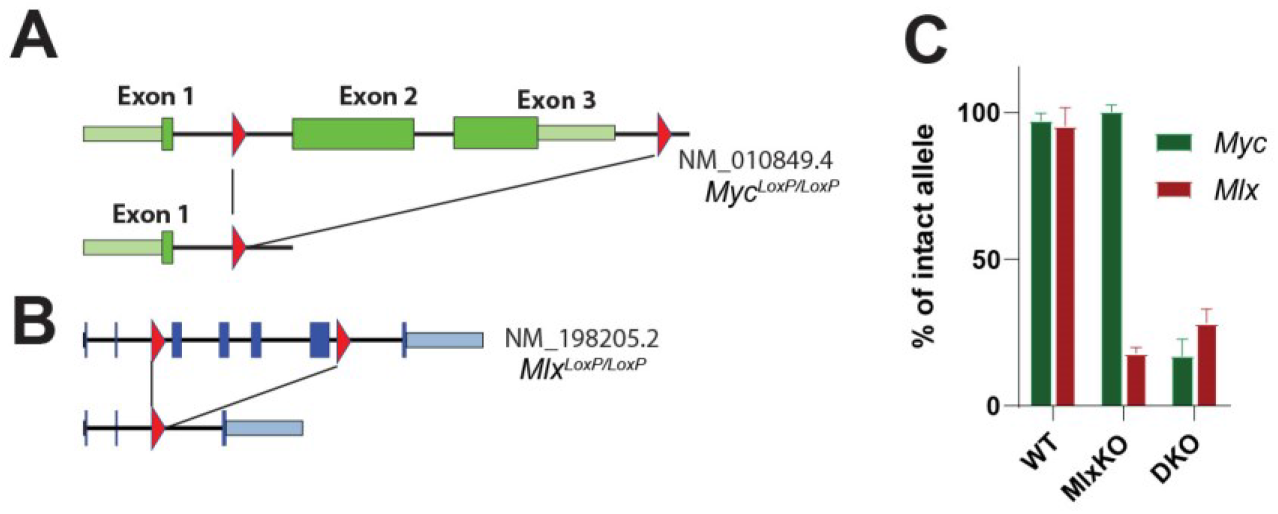
Strategies for the quantification of total donor and recipient hepatocytes and donor sub-populations. (A). Relevant regions of the murine *Myc* locus prior to and following CreER-mediated recombination showing the location of LoxP sites flanking coding exons 2 and 3 (red triangles) [25, 37]. (B). Relevant regions of the *Mlx* locus prior to and following CreER-mediated recombination showing the location of loxP sites flanking coding exons 3 and 6 [8]. (C) Verification of *MlxKO* and DKO knockouts. 4-5 wks after CreER activation, DNAs from the indicated livers were assessed for the presence of intact or recombined *Myc* and *Mlx* alleles as described in A-C. Hepatocytes were then used for transplant studies. Low-levels of non-excised genes likely originate from the non-hepatocyte populations present in the liver [25, 37].

Using FRG-NOD mice as recipients, we previously showed that WT donor hepatocytes outcompeted an equal number of *Chrebp*KO hepatocytes whereas WT and *Myc*KO hepatocytes competed equally [25, 35, 37]. Suspecting that *Mlx*KO hepatocytes would be even more defective, and to emphasize this, we delivered a total of 3×10^5^ donor hepatocytes intra-splenically into recipient mice at a ∼1:6 WT:*Mlx*KO ratio (Fig. 2A & B). After 24-28 wk of NTBC cycling, a number of recipients had died and no survivors had achieved NTBC independence, possibly as a result of the deliberate under-representation of WT hepatocytes in the initial inoculum. Indeed, quantification of the total donor population in the surviving recipients indicated that it comprised only 2-46% of all hepatocytes, which is both lower and more variable than typically achieved when mice receive larger numbers of replication-competent donor cells (Fig. 2C) [25, 37]. Despite this low level reconstitution, the surviving donor hepatocytes were nearly all WT despite their initial minority status (P<0.001) (Fig. 2D).

**Fig. 2.**
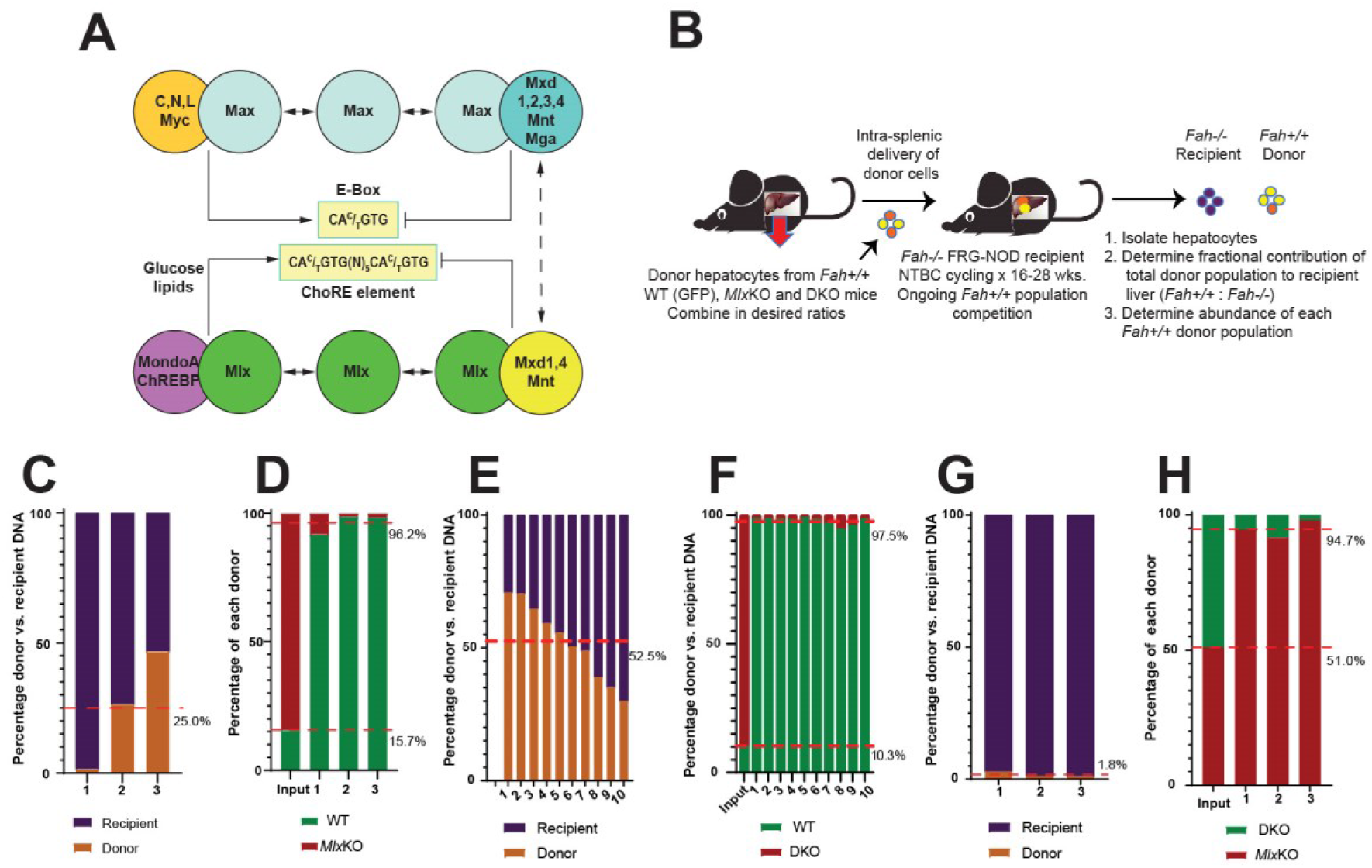
WT hepatocytes outcompete *Mlx*KO and DKO hepatocytes in repopulation assays. (A). The Extended Myc Network. The top portion of the panel shows the Myc Network, comprised of Myc, Max, Mxd1-4, Mnt and Mga1 and their consensus E-box binding site. The bottom portion shows the Mlx Network and its E-box-related but more complex ChoRE binding site [27, 46]. Mlx interacts with the nutrient-regulated positive factors ChREBP and MondoA or the negative factors Mxd1, Mxd4 and Mnt [2, 13, 24]. The latter cross-talk with the Myc Network (dotted arrow). (B). Hepatocyte transplantation strategy. Isolated *Fah+/+* WT or KO hepatocytes were mixed at the desired ratio and injected intrasplenically into FRG-NOD *Fah*^™/™^ mice maintained continuously on NTBC. NTBC cycling was continued until mice achieved NTBC independence or for 24-28 wks at which time total hepatocytes were isolated and the fractional representation of the total donor and recipient populations determined (Fig. 1). The contribution of each donor set was then further determined and compared to that of the input inoculum. (C). Following intrasplenic injection of 3×10^5^ donor hepatocytes comprised of a 1:6 ratio of WT and *MlxKO* cells, NTBC cycling was continued for 24-28 wks in the three animals that survived, with none achieving NTBC independence. Hepatocyte DNA was isolated from the transplanted animals and the percentage of recipient and total donor cells was determined. (D). DNAs from C were used to determine the ratio of the WT and *MlxKO* donor populations. DNA from an aliquot of hepatocytes at the time of transplant was used to confirm the input donor cell ratio. (E). Transplants performed in FRG-NOD *Fah*^™/™^ mice using inocula containing a ∼1:10 ratio of WT:DKO cells. Hepatocytes isolated after 24-28 wks of NTBC cycling showed that, on average, 52.5% of hepatocytes were comprised of donor cells. (F). Hepatocyte DNAs from E were used to determine the fraction of WT and *MlxKO* donor hepatocytes. Lane 1 shows the ∼1:10 ratio of the initial input inoculum. (G). Transplants performed in FRG-NOD *Fah*^™/™^ mice using inocula containing a ∼1:1 ratio of *MlxKO* and DKO cells. Total hepatocytes isolated after 24-28 wks of NTBC cycling were evaluated for the fractional representation of total donor and recipient populations showing that, on average, <2% of the total hepatocyte mass was of donor origin. (H). Fractional make-up of the donor population from G. Lane 1 shows the ∼1:1 ratio of the input population.

Although *Myc* deletion alone does not confer a replicative disadvantage to hepatocytes, *Chrebp* deletion does and is further exacerbated by the concurrent inactivation of *Myc* [25]. This suggests that the Myc and Mlx Networks are redundant and compensate for one another under certain circumstances. Because *Myc*KO x *Chrebp*KO hepatocytes still express MondoA [25], we asked whether its redundancy might mask more prominent phenotypes. We therefore compared the replicative potential of a mixed population of WT and DKO hepatocytes (1:10 ratio) in which the latter cells have functionally inactivated both ChREBP and MondoA as a consequence of *Mlx* deletion. This experiment achieved a somewhat greater rate of transplant success, with more animals surviving and with recipient livers eventually containing >50% donor hepatocytes (Fig. 2E). As before, however, virtually all these were of WT origin (P<0.001) (Fig. 2F).

To determine more directly which KO population was more proliferatively challenged, an additional competitive transplant experiment was performed using a 1:1 input ratio of *Mlx*KO and DKO donor hepatocytes. Overall survival was again low, no animals achieved NTBC-independence and <2% of hepatocytes isolated from recipients were of donor origin (Fig. 2G). However, despite their own inherent replicative compromise, *Mlx*KO hepatocytes showed an overwhelming survival advantage (Fig. 2H) and comprised nearly 95% of the recovered donor population (P<0.001). Together, these results argue that, in a highly demanding, long-term model of liver regeneration [25, 37, 46, 47], *Mlx* loss and the ensuing functional inactivation of ChREBP and MondoA markedly compromise donor hepatocyte proliferation and/or survival with the additional loss of *Myc* strongly reinforcing the defect.

### Overlapping transcriptional dysregulation in MlxKO and DKO livers primarily involves genes with roles in mitochondrial structure and function and translation

Prior to comparing whole transcriptome profiles of WT, *Mlx*KO and DKO livers, we confirmed that the latter two evidenced the predicted dysregulation of their direct target genes. For this, gene set enrichment analysis (GSEA) [48] was performed on three collections of direct Myc target genes from the Molecular Signatures Data Base C2 collection (MSigDB) (http://www.gsea-msigdb.org/gsea/msigdb/collections.jsp) and a 154 member panel of MondoA/ChREBP/Mlx direct target genes from the Qiagen Ingenuity Pathway Analysis (IPA) Data set (Table 2). In the first case, two of the three Myc target gene sets were also significantly enriched in *Mlx*KO liver RNA seq profiles, indicating as previously shown that some Myc-regulated genes are also responsive to Mlx Network inactivation (Fig. 3A) [13, 24, 25]. That the enrichment of these transcripts was broader and more pronounced in DKO livers, indicated that the Myc Network further contributes to the Mlx Network-mediated regulation of these targets as expected. In the second case, MondoA/ChREBP/Mlx target genes were significantly enriched in both *Mlx*KO and DKO livers (Fig. 3B). These results confirmed that *Myc* and/or *Mlx* inactivation was associated with both unique and shared responses of each Network’s respective target genes.

**Table 2.** MondoA, ChREBP and Mlx direct target genes from the Qiagen Ingenuity Pathway Analysis Data set.

| **Symbol** | **Entrez Gene ID** | **Symbol** | **Entrez Gene ID** | **Symbol** | **Entrez Gene ID** | **Symbol** | **Entrez Gene ID** |
| --- | --- | --- | --- | --- | --- | --- | --- |
| *Acaca* | 107476 | *Mrc1* | 17533 | *Rpl30* | 19946 | *Rps2* | 16898 |
| *Acacb* | 100705 | *Mtor* | 56717 | *Rpl31* | 114641 | *Rps20* | 67427 |
| *Acly* | 104112 | *Nr0b1* | 11614 | *Rpl32* | 19951 | *Rps21* | 66481 |
| *Acss2* | 60525 | *Nr1d1* | 217166 | *Rpl35* | 66489 | *Rps23* | 66475 |
| *Adgre1* | 13733 | *Pck1* | 18534 | *Rpl35a* | 57808 | *Rps24* | 20088 |
| *Adipor2* | 68465 | *Pgd* | 110208 | *Rpl36* | 54217 | *Rps25* | 75617 |
| *Agps* | 228061 | *Pklr* | 18770 | *Rpl36a* | 19982 | *Rps26* | 27370 |
| *Anxa2* | 12306 | *Plin1* | 103968 | *Rpl36al* | 66483 | *Rps27* | 57294 |
| *Arntl* | 11865 | *Pnpla2* | 66853 | *Rpl37* | 67281 | *Rps27rt* | 100043813 |
| *Ccl2* | 20296 | *Pnpla3* | 116939 | *Rpl37a* | 19981 | *Rps27a* | 78294 |
| *Ccl7* | 20306 | *Ppara* | 19013 | *Rpl38* | 67671 | *Rps27l* | 67941 |
| *Ccn3* | 18133 | *Pparg* | 19016 | *Rpl39* | 67248 | *Rps28* | 54127 |
| *Cebpa* | 12606 | *Ppargc1a* | 19017 | *Rpl39l* | 68172 | *Rps29* | 20090 |
| *Cidec* | 14311 | *Pygl* | 110095 | *Rpl3l* | 66211 | *Rps3* | 27050 |
| *Cpt1a* | 12894 | *Rgs16* | 19734 | *Rpl4* | 67891 | *Rps3a1* | 20091 |
| *Dhrs7b* | 216820 | *Rpl10* | 110954 | *Rpl41* | 67945 | *Rps4y1* | 20102 |
| *Elovl6* | 170439 | *Rpl10a* | 19896 | *Rpl5* | 100503670 | *Rps5* | 20103 |
| *Fabp4* | 11770 | *Rpl11* | 67025 | *Rpl6* | 19988 | *Rps6* | 20104 |
| *Fasn* | 14104 | *Rpl12* | 269261 | *Rpl6l* | 432502 | *Rps7* | 20115 |
| *Fgf21* | 56636 | *Rpl13* | 270106 | *Rpl7* | 19989 | *Rps8* | 20116 |
| *Foxa1* | 15375 | *Rpl13a* | 22121 | *Rpl7a* | 27176 | *Rps9* | 76846 |
| *Foxa2* | 15376 | *Rpl14* | 67115 | *Rpl7l1* | 66229 | *Rpsa* | 16785 |
| *G6pc* | 14377 | *Rpl15* | 66480 | *Rpl8* | 26961 | *Scap* | 235623 |
| *Gnpat* | 14712 | *Rpl17* | 319195 | *Rpl9* | 20005 | *Scd* | 20249 |
| *Gpam* | 14732 | *Rpl18* | 19899 | *Rplp0* | 11837 | *Sirt1* | 93759 |
| *Gpat3* | 231510 | *Rpl18a* | 76808 | *Rplp1* | 56040 | *Slc2a2* | 20526 |
| *Gpd1* | 14555 | *Rpl19* | 19921 | *Rplp2* | 67186 | *Slc2a4* | 20528 |
| *Hmgcr* | 15357 | *Rpl21* | 19933 | *Rps10* | 67097 | *Srebf1* | 20787 |
| *Hnf1a* | 21405 | *Rpl22* | 19934 | *Rps11* | 27207 | *Srsf2* | 20382 |
| *Hnf4a* | 15378 | *Rpl22l1* | 68028 | *Rps12* | 20042 | *Thbs2* | 21826 |
| *Igf2* | 16002 | *Rpl23* | 65019 | *Rps13* | 68052 | *Thrb* | 21834 |
| *Il1rn* | 16181 | *Rpl23a* | 268449 | *Rps14* | 20044 | *Thrsp* | 21835 |
| *Itgax* | 16411 | *Rpl24* | 68193 | *Rps15* | 20054 | *Tkfc* | 225913 |
| *Khk* | 16548 | *Rpl26* | 19941 | *Rps15a* | 267019 | *Tkt* | 21881 |
| *Lgals3bp* | 19039 | *Rpl27* | 19942 | *Rps16* | 20055 | *Txnip* | 56338 |
| *Lipe* | 16890 | *Rpl27a* | 26451 | *Rps17* | 20068 | *Ucp1* | 22227 |
| *Mlx* | 21428 | *Rpl28* | 19943 | *Rps18* | 20084 | *Ucp3* | 22229 |
| *Mlxip* | 208104 | *Rpl29* | 19944 | *Rps19* | 20085 |  |  |
| *Mlxipl* | 58805 | *Rpl3* | 27367 | *Rps19bp1* | 66538 |  |  |

**Fig. 3.**
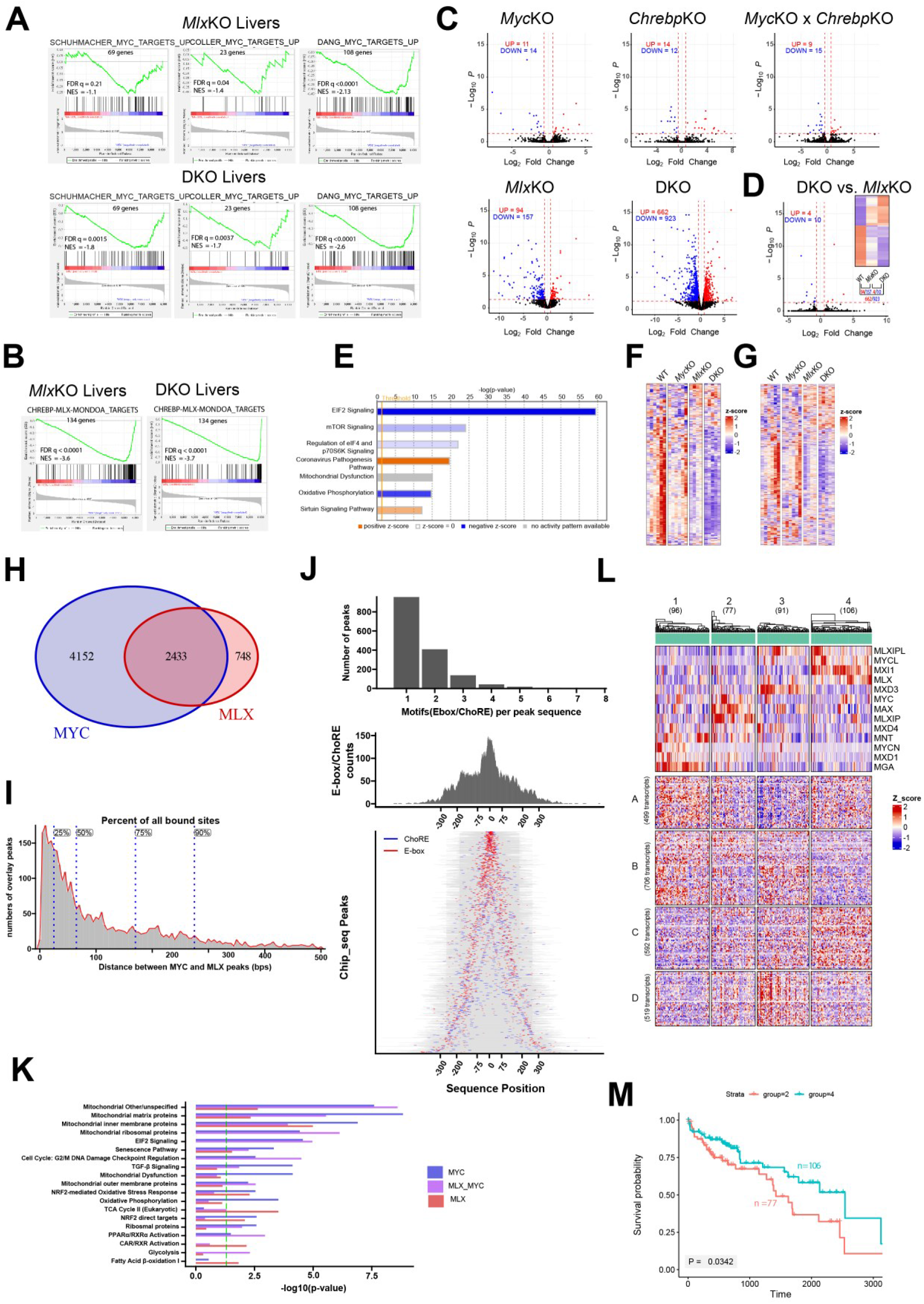
Transcriptional dysregulation in response to Myc and/or Mlx Network inactivation. (A). GSEA performed on RNAseq data sets obtained from *MlxKO* and DKO livers. Three sets of direct Myc target genes from the MSigDB Collection containing 69, 23 and 108 members, respectively, were used in the analysis, with expression levels being compared to those of WT livers (N=5 samples per group). (B). A 154 member collection of direct MondoA, ChREBP and Mlx target genes from the Qiagen IPA data set was used in GSEA on the samples shown in A (Table 2). (C). Volcano plots of differentially expressed genes in the indicated livers expressed relative to WT livers. Red and blue points = up-regulated and down-regulated, respectively relative to WT livers. *Chrebp*KO and *Myc*KO x *Chreb*KO liver RNA seq results were obtained from Wang et al. [25]. (D). Comparison of *MlxKO* and DKO livers demonstrating the differential expression of only 14 transcripts. The inset shows a heat map of differentially expressed transcripts among WT, *Mlx*KO and DKO livers. (E). IPA analysis showing the top seven dysregulated pathways in DKO livers. (F). Heat maps of expression differences for 260 transcripts from the MSigDB C2 data base encoding ribosomal subunits and proteins involved in translation (https://www.gsea-msigdb.org/gsea/msigdb/cards/REACTOME_TRANSLATION). (G). Expression differences for 605 transcripts encoding proteins comprising the mitochondrial proteome were compiled from the MitiProteome Data Base (http://www.mitoproteome.org). (H). Myc and Mlx binding sites in HepG2 cells. ChIP-seq results were downloaded from the ENCODE data base (https://www.encodeproject.org/) and analyzed for consensus Myc and Mlx binding sites residing within +/-2.5 kb of transcriptional start sites. The Venn diagram shows genes that bound only Myc, only Mlx or both factors. (I). Proximity of Myc and Mlx binding sites within the common target genes shown in H. Of the 2433 genes shown, a total of 5267 Myc and Mlx binding sites with overlapping footprints were identified. The positions corresponding to the peak center for each factor and the distances between them were determined. (J). The location and identities of E-boxes and ChoREs in relation to each factor’s binding site peaks. Top panel: the number of E-boxes and/or ChoREs associated with each footprint. Middle panel: the proximity of all motifs in relation to their site peak centers (designated as “0”). Bottom panel: The actual location of E-box and ChoRE motifs in each fragment and their position relative to the peak centers. Gray bars correspond to the length (in bps) of sequences that were determined. Some genes are depicted more than once as they contained more than a single non-overlapping binding site. (K). Select functional categories of genes represented by each of the three subsets of genes depicted in (H). Green dotted line: P < 0.05. (L). The transcriptomes (HTSeq-FPKM-UQ files) of 371 primary HCCs from TCGA were down-loaded by TCGAbiolinks R package [98] and then assigned to one of four groups (1-4) based on the expression of the indicated members of the Extended Myc Network determined using the ComplexHeatmap R package [99]. Tumors within these groups could be assigned to four additional categories (A-D) based on the expression patterns of the common 2433 transcripts shown in H. 116 of the genes are not shown as their expression was not reported in the TGCA database. (M). Comparative survival of individuals from groups 2 and 4 in (L).

To assess the effect of progressive dismantling of the Extended Myc Network on target gene sets, volcano plots were used to compare individual gene expression profiles in the above livers and previously described *Myc*KO, *Chrebp*KO and *Myc*KO x *Chrebp*KO livers (GEO Accession numbers GSE114634) [25]. In the latter three cases, <30 differences were identified relative to normal livers from age-matched animals (differential expression ≥1.5-fold and q<0.05) whereas *Mlx*KO and DKO livers showed up to 60-fold more differences (Fig. 3C). This more pronounced gene dysregulation again suggested that the combined loss of ChREBP and MondoA eliminated all redundant functions from the Mlx Network, thereby allowing a much larger complement of gene expression differences to be revealed. The relatively few differences between *Mlx*KO and DKO expression profiles (Fig. 3D and insert) was consistent with the notion that, at least in the normal, non-proliferating liver, the Mlx Network contributes more to regulating both the direct and indirect targets of both Networks. The differences between the DKO vs. *Mlx*KO groups in Fig. 3D were thus comparable to those between WT and *Myc*KO groups shown in Fig. 3C.

IPA profiling of the differentially expressed transcripts in DKO livers (and by extension *Mlx*KO livers) showed that six of the top seven most affected pathways were those with roles in mRNA translation and its control, energy metabolism and mitochondrial structure and function (Fig. 3E). The seeming exception (“Coronovirus Pathogenesis Pathway”) contained numerous ribosomal protein transcripts whose dysregulation accounted for this pathway’s inclusion. Collectively, these findings agreed with previous reports in livers, liver cancers and other cell types showing the Extended Myc Pathway’s role in the above processes [1, 2, 13, 17, 18, 24, 25, 37, 47, 49-52]. Gene expression profiles compiled from the pathways shown in Fig. 3E revealed the down-regulation of numerous transcripts encoding proteins involved in translation and mitochondrial structure and function in *Myc*KO and *Mlx*KO livers and an even greater degree of down-regulation in DKO livers (Fig. 3F&G) [25]).

To explore the potential co-regulation of direct target genes by the Myc and Mlx Networks, we obtained data from the most current version of the Encyclopedia of DNA Elements data base (ENCODE: https://www.encodeproject.org/) [53] and focused on the HepG2 HCC cell line, which was deemed the most relevant to the current work. CRISPR editing had inserted in-frame 3xFLAG epitope tags into the 3’-end of the endogenous *MYC* or *MLX* coding regions, thus allowing all ChIP-seq studies to be performed under identical conditions with the same anti-FLAG antibody. The results were trimmed using a default setting that enumerated only those binding sites within +/-2.5 kb of the transcriptional start site of each gene, thereby maximizing the likelihood of functional relevance. In this way, we identified 4152 genes that bound only Myc, 748 that bound only Mlx and 2433 that bound both factors at 6047 sites, 5267 of which overlapped, either entirely or partially (Fig. 3H). 37% of Myc target genes (2433/6558) also bound Mlx whereas 76% of Mlx target genes (2433/3181) also bound Myc, thus indicating that Mlx target genes are twice as likely to also bind Myc. 50% of the Myc and Mlx binding site peaks mapped to within 65 bp of one another and 75% mapped to within 170 bp, thus indicating that the two factors either bound to the same E-box or ChoRE or to more than one element in such close proximity that their individual peaks could not be resolved simply by examining the ChIP-seq footprints (Fig. 3I).

To confirm the above results and obtain greater resolution and characterization of Myc and Mlx binding, we analyzed the sequences flanking the sites of maximum factor binding (i.e. the ChiPseq binding “peaks”). The above 6047 binding sites were merged to 2863 distinct sites for motif analysis. 1220 of these (42.6%) contained consensus E-boxes for Myc-Max and 714 contained consensus ChoREs [54](Fig. 3J). 45.2% of the sites contained neither E-boxes nor ChoREs, despite the presence of prominent Myc and/or Mlx footprints, thereby indicating either that the motifs did not conform to the conservative consensus sequences used in our search or that Myc and Mlx binding was indirect as a result of association with other DNA-binding factors.

Remarkably, the E-boxes and ChoREs depicted in Fig. 3J displayed a non-random distribution and tended to occurr within close proximity of factor-binding peaks. The consensus binding sites located closest to or at the peak centers tended to be those whose adjacent sequences contained either the fewest numbers of additional motifs or tightly clustered ones.. This suggested that many ChIP-seq peaks represent the integrated signal of multiple variably overlapping and unresolvable individual binding sites and thus do not necessarily directly overlie a particular site. Collectively, these findings confirm the presence of multiple E-boxes and/or ChoREs within the majority of common target genes as well as direct evidence of Myc’s binding to ChoREs and Mlx’s binding to E-boxes in select cases. They further suggest a means by which the apposition of multiple binding sites within some genes could allow for the simultaneous binding of different combinations of factors as well as their direct interaction and cross-talk.

Cataloging the functions of the 2433 common genes shown in Fig. 3H using the IPA and Mitoproteome data bases and a bespoke collection of previously published genes [25, 49, 55, 56] showed that many could be categorized as supporting mitochondrial and ribosomal structure and function (Fig. 3K) [25, 55]. Thus, the common Myc- and Mlx-bound target genes in human HepG2 cells faithfully reflect the both the current transcript changes and those previously documented in *Myc*KO and/or *Chreb*KO murine livers and hepatoblastomas [25, 55]. Interestingly, the 4152 genes bound only by Myc and the 748 genes bound only by Mlx fell into somewhat different IPA categories than did the common genes (Fig. 3K). For example, Myc-specific genes were also involved in more restricted and/or unique functions such as TGF-β signaling, cell cycle and oxidative phosphorylation Similarly, Mlx-specific genes also tended to belong to distinct functional categories such as those related to retinoic acid signaling, xenobiotic metabolism, fatty acid β- oxidation and the TCA cycle.

Given the Extended Myc Network’s dynamic nature (Fig. 2A), the potential for different members to bind multiple closely neighboring sites with different affinities (Fig. 3J) and their ability to either augment or antagonize one another’s transcriptional impact, we hypothesized that Myc and Mlx binding alone (Fig. 3I and J), would not necessarily predict target gene expression levels. We further hypothesized that the transcriptional impact on any individual target gene would ultimately reflect the entire Extended Network’s integrated action [57]. We thus examined the expression of the 2433 common Myc and Mlx target genes (Fig. 3H) in 371 human HCCs using data from The Cancer Genome Atlas (TCGA). Target gene expression could be categorized into four groups (designated A-D) that correlated with four patterns of Extended Myc Network member expression (groups 1-4) (Fig. 3L). Two of the tumor groups also showed significant differences in survival (Fig. 3M). Together with the results of Fig. 3J, these findings support the idea that the binding of Myc, Mlx or any other Extended Myc Network factor to a target gene likely reflects only one aspect of the complex and integrative interplay among other Network members that collectively dictates the gene’s expression level and downstream biological consequences[20].

Myc, particularly when it is over-expressed by tumor cells, promotes the Warburg effect by up-regulating glucose transporters and genes encoding glycolytic enzymes [12, 25, 40, 51, 58-63]. However, none of these showed altered expression in *Myc*KO livers (Table 3). However, the progressive inactivation of the Extended Myc Network was associated with the down-regulation of three transcripts, all of which encode rate-limiting transporters or enzymes. These included the major hepatocyte glucose transporter Glut2/Slc2a2 and the glycolytic enzymes liver-type phosphofructokinase (Pfkl) and liver type pyruvate kinase (Pklr). Glut2/Slc2a2 is also required for the proper regulation of glucose-sensitive genes and for glucose-stimulated insulin secretion [62, 64]. These findings were consistent with the previous IPA showing that genes comprising a glycolysis-related set were co-bound by Myc and Mlx (Fig. 3K).

**Table 3.**
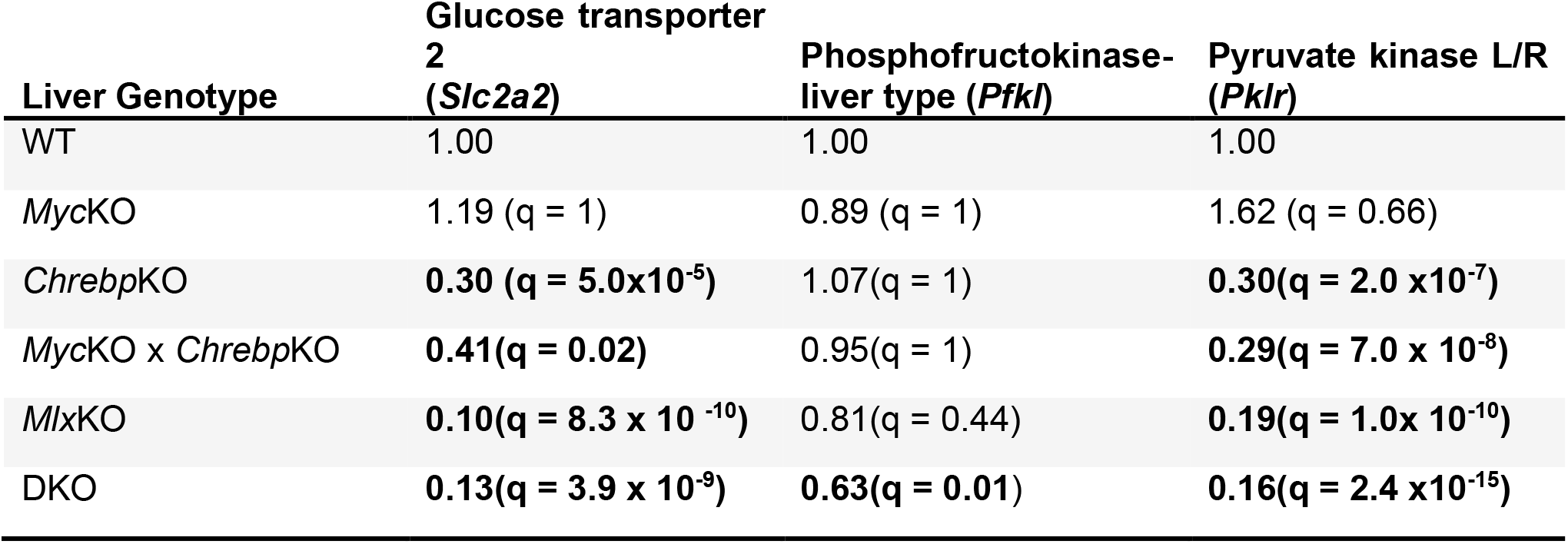
Relative expression levels of transcripts for rate-limiting factors in the glycolytic pathway.

### Loss of the Extended Myc Network members causes NAFLD

Consistent with findings that the Myc and Mlx Networks both impact pathways involved in carbohydrate and lipid metabolism (Fig. 3K) [1, 2, 7, 24, 25, 37, 65, 66], young *Myc*KO, *Chrebp*KO and *Myc*KO x *Chrebp*KO mice develop NAFLD [25, 37, 67]. However, these studies did not determine if this was progressive or if the dual compromise of the Myc and Mlx Networks increased its severity. We therefore examined the livers of older (14-16 month) *Myc*KO, *Chrebp*KO, *Myc*KO x *Chrebp*KO, *Mlx*KO and DKO mice to evaluate the extent of NAFLD. Relative to WT livers, all KO livers showed more intense Oil Red O staining but did not significantly differ from one another (Fig. 4A-F). They also contained more total triglyceride than did livers from younger *Myc*KO, *Chrebp*KO and *Myc*KO x *Chrebp*KO mice [25, 37] (Fig. 4G). These findings suggest that NAFLD appears earlier in *Mlx*KO and DKO mice but that *Myc*KO mice eventually achieve a similar degree of severity.

**Fig. 4.**
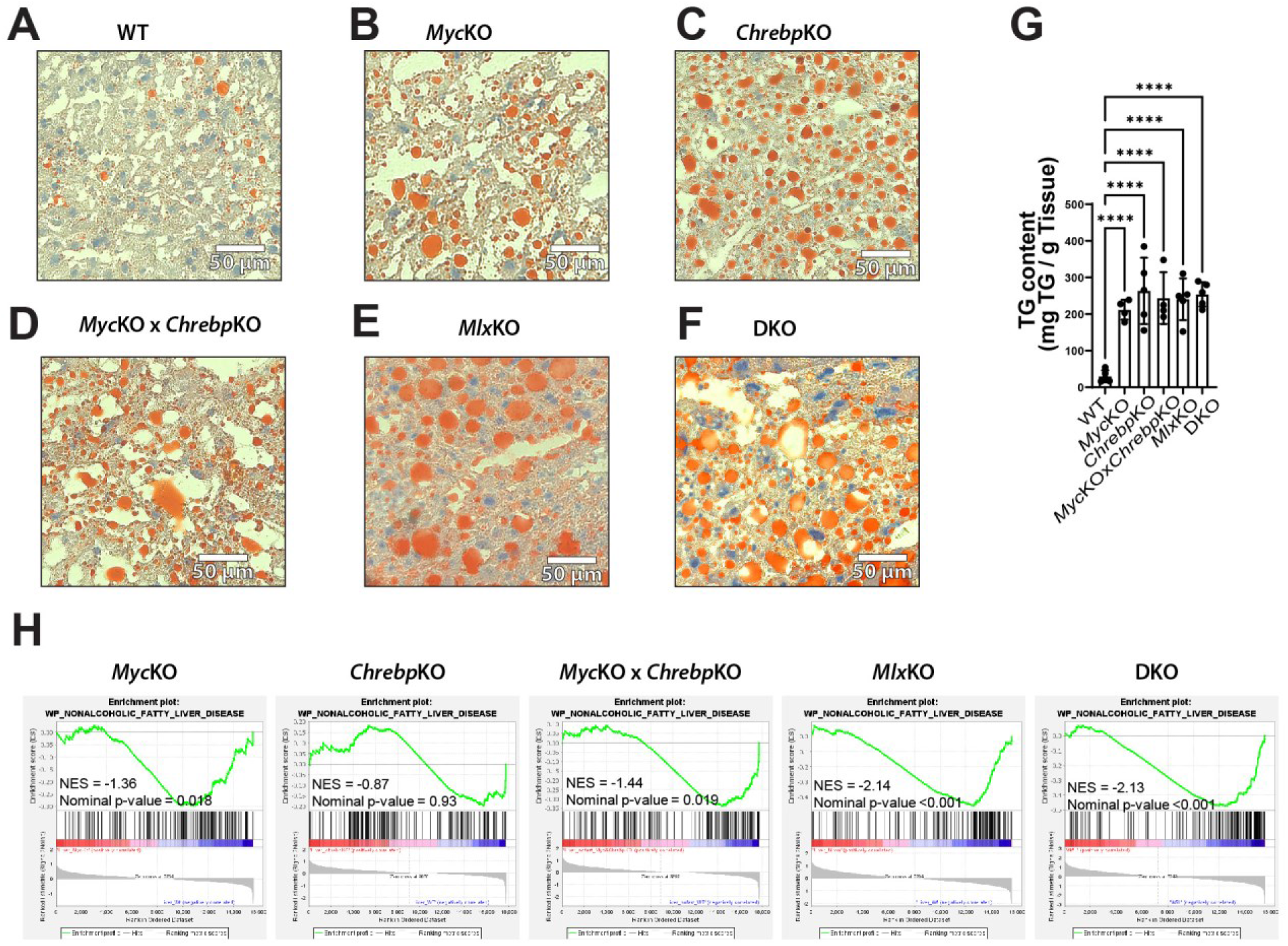
Neutral lipid accumulation is a feature of livers with compromise of the Extended Myc Network. (A-F). ORO stained sections of livers from 14-16 mo mice of the indicated genotypes. (G). Quantification of triglyceride levels in liver samples from A-F. (H). Enrichment of genes involved in human NAFLD. RNAseq results from the indicated groups of KO mice were compared to their age-matched WT counterparts for the expression of a gene set involved in NAFLD (https://www.wikipathways.org/instance/WP4396_r98945). Normalized enrichment scores (NES) and nominal P values are shown above each profile.

In further support of the above conclusions, we also found evidence for enrichment of a 163 member gene set associated with human NAFLD in all cohorts except *Chrebp*KO, (https://www.wikipathways.org/instance/WP4396_r98945) (Fig. 4H). Thus, despite the fact that the causes of NAFLD in the above mice and humans differ considerably, there is in most cases considerable similarity in the disease-related gene expression profiles [68].

Finally, we performed IPA to identify additional disease-related pathways that are sometimes dysregulated in NAFLD [68] and in all cases found several relating to lipid synthesis/metabolism and PPAR activation (Table 4). While noting little histologic evidence for inflammatory cell infiltrates or fibrosis in our KO livers, several of these pathways were in fact associated with non-alcoholic steatohepatitis (NASH) (Table 4). Although these transcriptome-based findings may represent early evidence of actual NASH, the altered expression of inflammatory markers could also be indicative of the immune function changes that accompanies Myc dysregulation in non-hepatic tissues [37, 69]. We believe the most conservative interpretation of the above results is that the NAFLD that accompanies the loss of most Extended Myc Network members exhibits evidence of mild NASH as indicated by significantly altered molecular markers of this state but little documentable histo-pathological change.

**Table 4.**
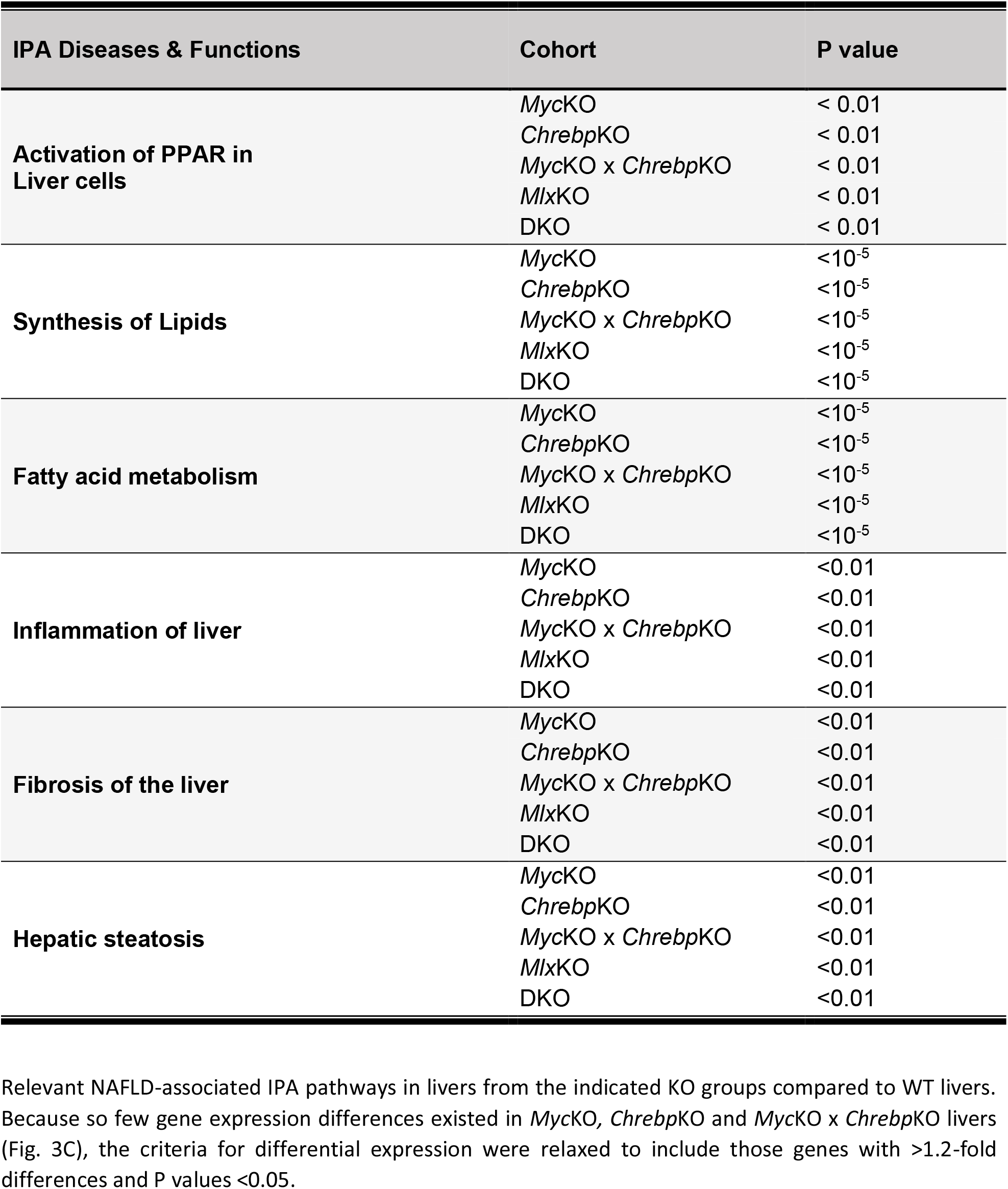
IPA profiling of KO livers.

### MlxKO and DKO mice develop age-related hepatic adenomatosis and occasional HCC

Unexpectedly, 36% of *Mxl*KO and DKO animals (15 of 42) of both genders developed multiple small-medium-sized hepatic neoplasms, which were never observed in WT, *Myc*KO, *Chrebp*KO or *Myc*KO *x Chrebp*KO mice (Fig. 5A&B) [25, 37]. These were mostly well-differentiated and/or myxoid-type tumors with numerous balloon cells, nuclear enlargement and microvesicular steatosis. A minority also showed small foci of well-differentiated HCC, which is sometimes associated with hepatic adenomas in humans (Fig. 5C) [44, 70]. Regardless of histology, adenomas showed significantly more staining for Ki-67 than did the adjacent non-neoplastic liver parenchyma (Fig. 5D, P=3.4×10^-5^).

**Fig. 5.**
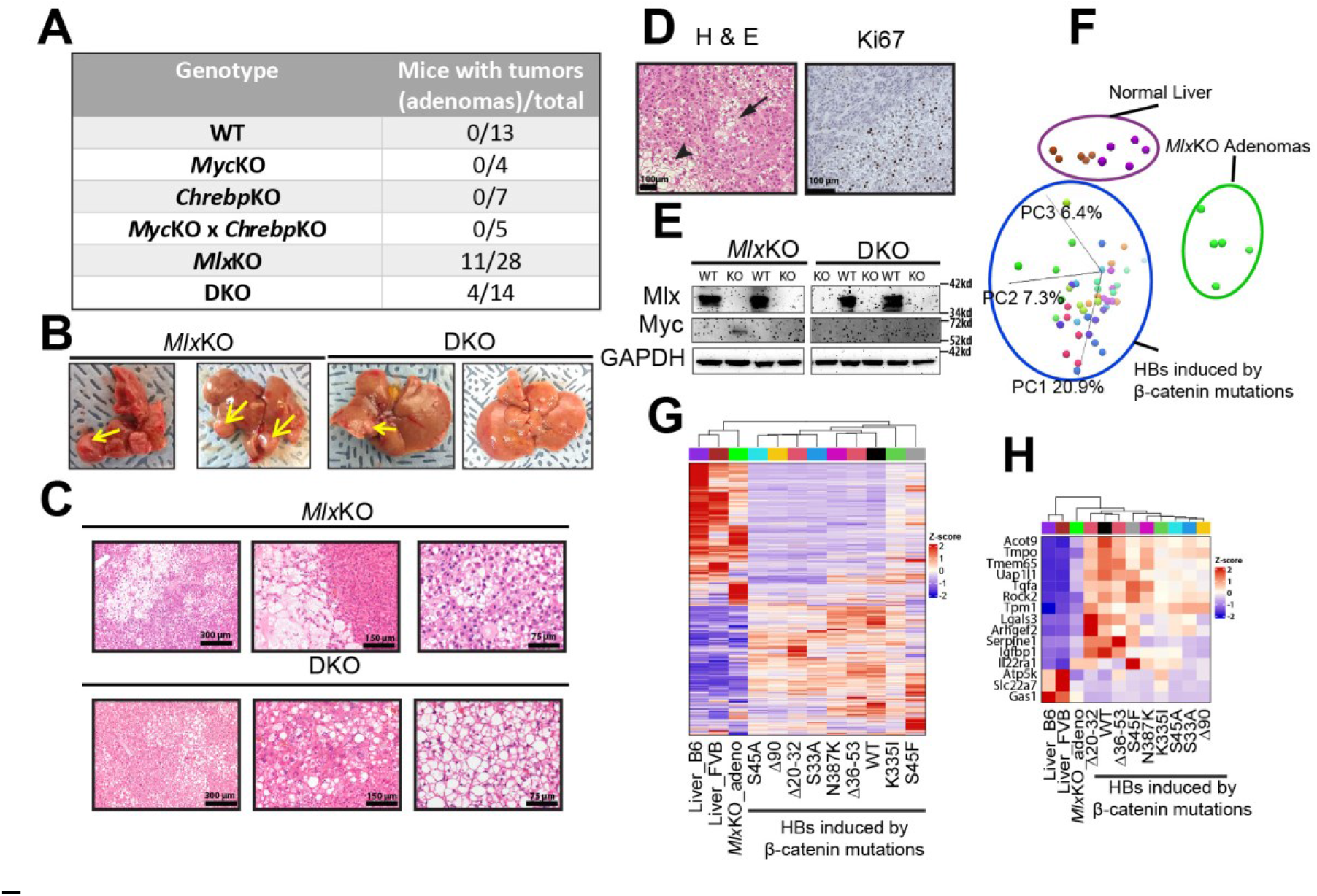
Characterization of hepatic adenomas originating in *MlxKO* and DKO livers. (A). Number of 14-16 mo mice of the indicated genotypes with visible liver tumors at the time of sacrifice. (B). Gross appearance of tumor-containing livers from A. Tumors were generally small but multifocal (arrows). (C). H&E-stained sections showing the typical appearance of *MlxKO* and DKO livers. The first two *MlxKO* images depict well-differentiated and myxoid-type adenomas, respectively, whereas the third panel depicts a focus of well-differentiated HCC embedded within an adenoma. The first two DKO images depict regions of microvesicular steatosis with balloon cells and nuclear enlargement, respectively whereas the third image shows another adenoma that is quite similar in appearance to those arising in *MlxKO* livers. (D). H&E and Ki-67 immuno-stained sections of the Myc-expressing adenoma from D showing regions of inflammation (arrowhead), adjacent to those resembling well-differentiated HCC (arrow). Ki-67 staining is more intense within the nodular adenoma (lower right) compared to adjacent normal liver (upper left). Quantification of Ki-67 staining from four different adenomas and adjacent normal tissue was performed on 300-2000 cells from each region. The mean Ki-67 index in adenomas was 30.08+/-6.5%% versus 3.2+/-3.1% in adjacent non-adnomatous tissues (P=3.4×10^-5^). (E). Immuno-blots showing Mlx expression in WT livers and its absence in adenomas arising from *MlxKO* and DKO livers. Only one large adenoma from an *MlxKO* mouse with hepatomegaly expressed detectable levels of Myc (lane 2). (F). PCA of whole transcriptomes from WT livers, HBs generated by the over-expression of mutant forms of β-catenin+YAP^S127A^ and the above-described adenomas. (G). Whole transcriptome profiles of the tissues from F. Note that control liver samples are derived from two different strains, C57B6 and FVB. (H). Heat maps of 15 of the 22 transcripts that are dysregulated in murine HBs and correlate with poor outcomes in human HBs and other human cancers [47].

Adenomas did not express Mlx protein, indicating that they did not originate from residual hepatocytes that had escaped *Mlx* locus excision and maintained their growth advantage (Fig. 5E). Normal livers and DKO adenomas also did not express detectable Myc protein nor did most *Mlx*KO adenomas. This was consistent with their slow growth rates and a likely consequence of their Extended Myc Network defects (Fig. 3C-G) [25, 37]. An exception was seen in a single large adenoma containing elements of HCC from an *Mlx*KO mouse with marked hepatomegaly (liver weight 6.7 g or approx. three times normal) (Fig. 5E).

We performed RNAseq on five adenomas from *Mlx*KO mice and compared their transcriptome profiles to those of WT livers and 45 primary murine hepatoblastomas (HBs) generated by over-expressing the Hippo pathway terminal effector YAP^S127A^ together with one of nine different patient-derived oncogenic mutations of β-catenin [71]. The transcriptional profiles of the adenomas were distinct from those of both livers and all HBs (Fig. 5F&G). Adenomas did not over-express wild-type β-catenin or YAP as is common for HBs [72] but did dysregulate 15 of 22 transcripts that are aberrantly expressed in all murine HBs regardless of etiology and that correlate with survival in human HB and other cancers (Fig. 5H) [47, 73]. Our results indicate that dismantling the Mlx Network, either alone or concurrently with Myc leads to the eventual emergence of multiple adenoma-like hepatic neoplasms (adenomatosis), which, like their human counterparts, can further evolve and acquire HCC-like features [70].

## Discussion

Most previous investigations into Myc’s role in hepatic regeneration have relied upon the PH model and yielded conflicting results that likely reflected differences in how and when regeneration was assessed and quantified [37]. The short time frame over which this process occurs and the dependency upon separate groups of mice may have further contributed to disparate outcomes. Because post-PH hepatocytes require fewer than two divisions to replace the missing mass, the model also poses a comparatively modest regenerative challenge. Indeed, even this low number overestimates the actual contribution made by dividing hepatocytes given that about half the response to PH involves hypertrophy of the liver remnant plus replicative contributions by non-hepatocyte populations such as endothelial, Kupfer and stellate cells [38, 74, 75]. In contrast, the FAH model is associated with a more sustained and robust 50-100-fold expansion of pure populations of transplanted hepatocytes. It also provides a well-defined point at which a stable level of regeneration can be assessed and a means by which competing donor populations within the same liver can be simultaneously distinguished and quantified following their delivery at any desired and pre-selected ratio [25, 37, 50]. Using this approach, we previously showed that the long-term proliferation of otherwise normal hepatocytes requires ChREBP but not Myc, although the loss of both factors was additive [25, 37]. Even more pronounced inter-dependencies were seen during HB tumorigenesis, with HB growth being markedly impaired in both *Myc*KO and *Chrebp*KO livers and even more so in *Myc*KO x *Chrebp*KO livers. [25, 46, 47], These findings implied a means of communication between the Myc and Mlx Networks, with each one being able to at least partially rescue defects in the other [1]. They also demonstrated that the requirement for Myc becomes progressively more critical as proliferative demand increases, thus emphasizing it strong contextual dependency (Fig. 6).

**Fig. 6.**
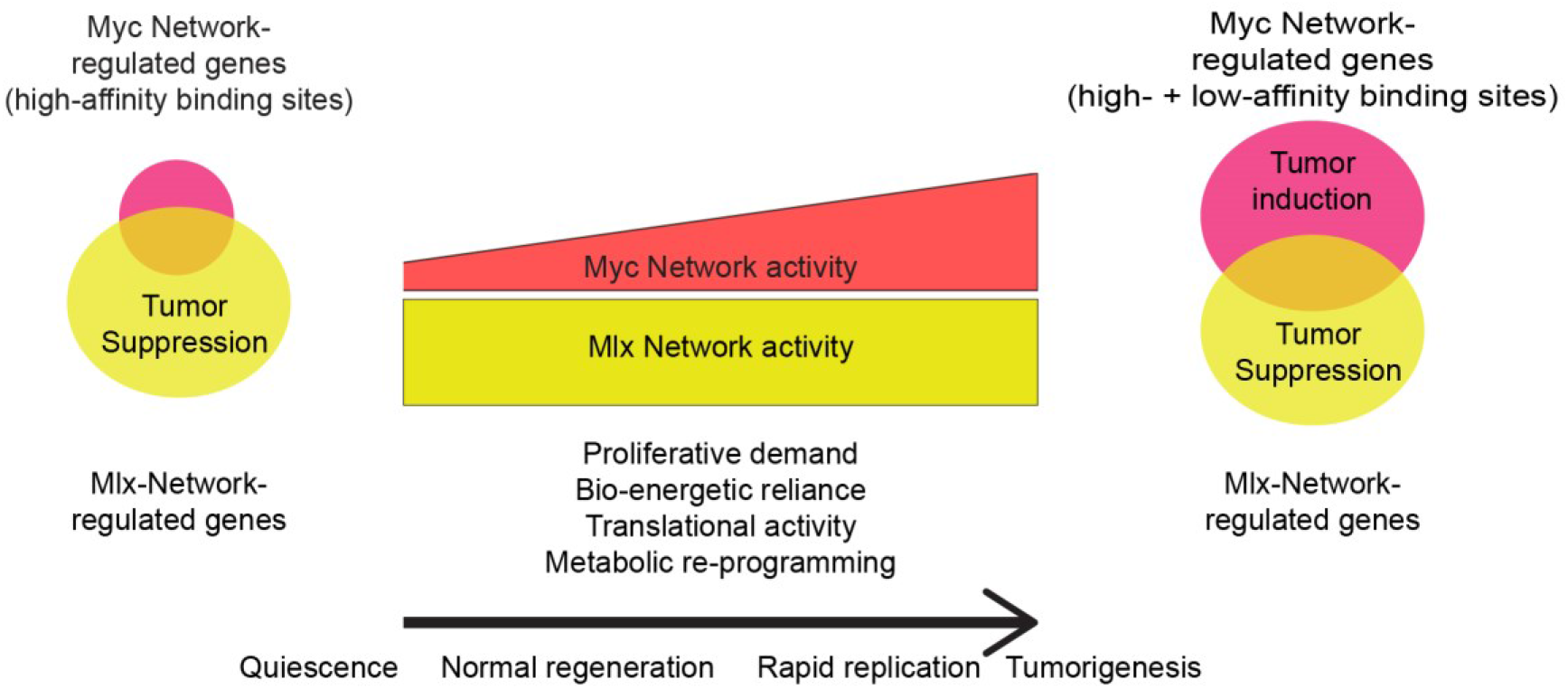
A model for gene regulation and control of normal and neoplastic proliferation by the Extended Myc Network. During quiescence, Myc levels are low and its target genes, which are relatively few in number (Fig. 3 and [76]) tend to be those with high-affinity E-box binding sites. Slow and controlled replication, such as that occurring during the replacement of *Fah-/-*hepatocytes, is largely regulated via the Mlx Network [25, 37]. As replication increases or as cells acquire transformed features, increased Myc expression [25, 49] activates genes with low-affinity binding sites, including those encoding glycolytic enzymes that are collectively responsible for the Warburg effect as well as other metabolic pathways and functions that support increased energy demands and rapid growth [2, 3, 24, 40, 51, 63, 67, 100]. Despite the rapid growth that occurs in response to the over-expression of Myc and mutant forms of β-catenin and YAP^S127A^ [25, 49, 71] the Mlx Network, which supports this rapid growth, is proposed to also contribute to tumor suppression as well (Fig. 5).

Despite our previous transplant studies having been performed with different input ratios of WT and KO hepatocytes [25, 37], their outcomes were consistent with those reported here and allowed us to extend our conclusions regarding the relative importance of the Myc and Mlx Networks in liver regeneration. For example, our studies comparing WT and *Chrebp*KO hepatocytes employed an input inoculum comprised of 62% of the latter population that was reduced by more than half following competitive repopulation [25]. Our current results in which *Mlx*KO hepatocytes comprised ∼84% of donor cells but only ∼4% of the final population (Fig. 2D) provided strong evidence that the concurrent functional inactivation of both ChREBP and MondoA confers an even more profound proliferative disadvantage. This could be a direct effect resulting from the concurrent loss of ChREBP and MondoA binding to their respective target genes either individually or collaboratively with Myc thereby eliminating any possibility of rescue of one factor by another (Fig. 2A) [24, 26-29]. A non-mutually exclusive indirect effect that allowed Mxd1, Mxd4 and Mnt to suppress Myc target genes more effectively by increasing their association with Max also remains possible (Fig. 2A). The relative importance of these two models could vary among different target genes, at different times during repopulation or in different liver compartments.

We also previously showed that the combined loss of *Myc* and *ChREBP* suppressed regeneration more than did the knockout of either individual gene, thereby corroborating previous evidence for inter-Network cross-talk [2, 3, 13, 24, 25]. In that study, performed with nearly equal contributions of WT and *Myc*KO x *Chrebp*KO donor hepatocytes, the latter was reduced to 7.5% following repopulation [25]. Though markedly impaired, the residual proliferative activity of these cells could have reflected the redundant function of MondoA (Fig. 2A), which is supported by two separate aspects of the current study. The first was the ∼45-fold repopulation advantage of WT hepatocytes over DKO hepatocytes, whereas the second was the ∼10-fold repopulation advantage of *Mlx*KO hepatocytes over DKO hepatocytes (Fig. 2F&H). Collectively our current results indicate that both the Myc and Mlx Networks play distinct as well as redundant roles in normal hepatocyte replication. However much of the proliferative drive needed to sustain hepatocyte expansion in FAH mice is subsumed by the Mlx Network regardless of the Myc Network’s status [25, 37]. This is supported by the progressive deterioration of repopulation potential as the Extended Network is gradually dismantled (Fig. 2C-H) [25].

In proliferatively quiescent cells or organs such as the liver, Myc is usually expressed at low levels and regulates relatively few genes in contrast to Mlx (Figs. 3C and 5E) [25, 37, 76]. Myc’s contribution to genome-wide transcription may therefore be better appreciated in tumors where its over-expression can activate genes that are otherwise non-physiologic targets due to their low-affinity binding sites [10, 11, 25, 66]. Another plausible explanation for the seemingly modest transcriptional consequences of Myc loss in some normal tissues is that at least some Myc target gene expression is maintained by the Mlx Network with redundant contributions being made by MondoA and/or ChREBP [25, 37, 76]. This is best appreciated in livers and tumors when the Myc and Mlx Networks are either individually and concurrently inactivated (Fig. 3C) [25].

In tumors, the Myc Network positively regulates most of the genes encoding glycolytic enzymes and strongly contributes to the Warburg effect [5, 12, 40, 49, 51, 59, 63, 77] as it does in rapidly growing fibroblasts *in vitro* [40, 77]. In contrast, our transcriptomic studies have not revealed such wide-spread roles for the Myc and Mlx Networks in maintaining glycolysis *in vivo* (Table 3 and Fig. 3K), which may reflect Myc’s low-level expression, the relative proliferative quiescence of the normal liver and its greater reliance on fatty acid oxidation as an energy source [25, 46, 49, 71]. Nonetheless, among the most down-regulated genes in *Mlx*KO and/or DKO livers, were *Glut2/Slc2a2, Pfkl* and *Pklr* whose encoded proteins are rate-limiting for glucose uptake and glycolysis. In rat insulinoma cells, the *pklr* proximal promoter binds both ChREBP and Myc, with the former interacting with a ChoRE element and the latter binding elsewhere [28, 29]. These results suggest that, in normal liver, glucose uptake and oxidation are more reliant on the Mlx Network (Fig. 6) whereas, in response to Myc-driven transformation or normal proliferation, more extensive transcriptional regulation of glucose uptake and its oxidation is achievable [4, 14, 23, 25, 49, 78]. This could have the additional benefit of maximizing glycolytic efficiency and sustaining cell division when micro-environmental glucose and oxygen supplies were limiting and nutrient-dependent functions of MondoA and ChREBP were attenuated [79].

Coordinated changes in direct and/or functionally-related Myc and Mlx Network target genes were identified in *Mlx*KO and DKO livers but were more pronounced in the latter (Fig. 3A-D) [25, 37]. Some have been previously shown to support protein translation and its control as well as mitochondrial structure and function (Fig. 3E-G) [18, 25, 37, 40, 49, 67, 71]. As was seen for individual glycolysis-related transcripts (Table 3), the collective expression of these sets became increasingly compromised as the Extended Myc Network was progressively inactivated. The dramatic up-regulation of these pathways that accompanies tumorigenesis in WT livers had also been shown to be attenuated in response to Myc and/or ChREBP inactivation and to correlate with slower tumor growth rates [25]. We provided a mechanistic underpinning for the coordinated response of the relevant gene sets associated with these pathways by showing that, in HepG2 cells, 37% of Myc’s direct target genes also bind Mlx while 76% of direct Mlx target genes also bind Myc (Fig. 3H). Many of the previously mapped binding sites for these two factors overlapped and/or contained multiple E-boxes and/or ChoREs (Fig. 3I and J). While this sometimes made it difficult to attribute precisely Myc or Mlx binding to a specific motif within a factor’s ChIPseq footprint, the collective binding landscape suggested a model for target gene regulation that accommodated this and all other observations. The model accounted for the fact that many sites containing only E-boxes coincided with Mlx binding peaks whereas many ChoRE-only sites coincided with Myc binding peaks (Fig. 3J). This indicated that cross-talk between the Myc and Mlx Networks occurs by virtue of shared common binding sites as previously suggested. The non-random distribution of E-boxes and ChoREs around Myc and Mlx binding peaks (Fig. 3J) also suggested that more than one such site could be occupied at any given time, that binding might be cooperative and that the composition of the bound factors, their interactions with each other and differential protein-DNA affinities serve to fine tune the target gene’s transcriptional output. While we examined only Myc and Mlx binding, these motifs could also bind other Extended Myc Network members such as those between Max and Mxd proteins, which would not have been detected with our ChIPseq analysis. Whether closely-spaced Mlx sites contained ChREBP or MondoA heterodimers could also potentially determine if, when and the degree to which a gene was responsive to metabolic substrate-mediated regulation. Finally, the four groups into which the expression patterns of the 2433 common Myc + Mlx direct target genes in HCCs could be compiled correlated with the patterns of Extended Myc Network transcript expression and, in two cases, with significant survival differences (Fig. 3L and M). In future studies, it will be important to determine the degree to which different neighboring heterodimeric combinations of Extended Myc Network members either cooperate with or antagonize one another under different conditions in different cell types.

Myc and/or ChREBP inhibition are widely associated with lipid accumulation, which stems from an over-reliance on fatty acid oxidation and a resulting increase in lipid uptake that exceeds the amount necessary to satisfy energy demands (Fig. 4) [25, 37, 65, 80, 81]. Young mice with hepatocyte-specific loss *of Chrebp* or combined *Myc + Chrebp* loss also accumulate more neutral lipid than do those with isolated Myc knockout [25]. While we did not serially follow these animals, our findings suggest that, early in life, the partial or complete inactivation of the Mlx Network promotes a more rapid genesis of NAFLD than does inactivation of Myc alone. Over time, however, lipid accumulation equalizes, with little differences among the various KO groups being discernible (Fig. 4G). KO livers dysregulated many of the same gene sets and/or IPA pathways that have been described in NAFLD and its progression in humans (Fig. 4H and Table 4) thereby further supporting the mechanistic relatedness of the various factors responsible for this state. Because KO livers also displayed molecular evidence of incipient NASH, it will be important in future work to determine whether these features become more pronounced with age and whether histological findings of inflammation and fibrosis become discernible [82, 83].

An unanticipated finding was the development of hepatic adenomatosis in more than one-third of *Mlx*KO and DKO mice (Fig. 5A&B). This incidence likely represents an underestimate since animals older than 14-16 mos. were not investigated and microscopic adenomas may have been overlooked in some instances. That similar neoplasms did not appear in WT, *Myc*KO, *Chrebp*KO or *Myc*KO x *Chrebp*KO mice makes it likely that complete Mlx Network inactivation is a pre-requisite for their development. While these neoplasms bear the hallmarks of actual human adenomas [44, 70], several features suggest more aggressive and malignant predilections, despite the lack of Myc expression in most. These include their multi-focality, their occasional HCC-like features and their robust Ki-67 expression (Fig. 5C&D). In contrast, human adenomas, while well-known for their occasional conversion to HCC, are typically few in number and tend to display only modestly higher Ki-67 expression [70]. Molecular features suggestive of more aggressive behavior in our adenomas include the dysregulation of 15 of 22 transcripts that we recently identified as predicting inferior outcomes in human HBs and over a dozen other human cancer types (Fig. 5H) [47].

Recurrent *MLX* gene deletions are associated with at least eight human cancer types and provide further reason to implicate the Mlx Network in the pathogenesis of hepatic adenomatosis (https://portal.gdc.cancer.gov/genes/ENSG00000108788[20]. Genetic suppressors of hepatic adenomas and other benign tumors such as meningiomas, neurofibromas and uterine fibroids are well-documented but are distinct from more notorious counterparts such as *TP53, RB, PTEN, APC* and *BRCA1*/2 that are associated with malignant tumors [84-87]. However, the role of Mlx and its members may also be more indirect and nuanced. For example, NAFLD (Fig. 4) is a known predisposing factor for the development of both adenomas and HCC and we are currently unable to determine how it might affect tumorgenesis in *Mlx*KO mice [70, 88, 89]. On the other hand, the failure of *Myc*KO, *Chrebp*KO or *Myc*KO *x Chrebp*KO mice to develop adenomas or HCCs, despite their equally pronounced NAFLD as well as the fact that high-fat diets can actually suppress hepatic tumor growth [90], argues for a more direct role for the Mlx Network in adenoma suppression. It will be of interest to determine whether *Mlx*KO mice are more susceptible to transformation by other oncogenic stimuli even though the emergence of the ensuing tumors may be delayed and their growth slowed.

In summary, we have shown that the Mlx Network engages in considerable biological and molecular cross-talk with the Myc Network and plays a more substantive role in long-term liver regeneration [25, 29, 35, 37, 46]. Both networks, but the former in particular, alter the expression of numerous genes responsible for broad aspects of translation and energy generation by both aerobic and anaerobic pathways [25, 37, 55]. The majority of Myc and Mlx targets are co-regulated or at least bound by both factors, which appear to share many of the same binding sites, which often lie in close proximity to one another. The actual expression of these targets genes further correlates with the patterns of expression of all 13 members of the Extended Myc Network thereby suggesting complex interactions and inter-dependent cross-talk at their sites of binding. Mechanistically, the defects that ensue in KO cells as a result of compromising these genes reflect an inability to maintain energy production and translation at levels commensurate with their proliferative demands. This is particularly acute in the neoplastic setting where tumor growth, but not induction, may be severely compromised [25, 37, 55]. The presumptive energy dysequilibrium that arises as a consequence of perturbing either or both of the networks is likely addressed by an increased uptake, oxidation and storage of fatty acids leading to eventual NAFLD [25, 71]. The hepatic adenomatosis and occasional HCC seen in response to Mlx Network compromise suggests that the tumor suppressor-like activity of the Mlx Network counters the more pro-oncogenic tendencies of Myc over-expression. Our findings emphasize the elaborate orchestration of the Extended Myc Network in balancing energy demands and metabolism with normal and neoplastic proliferation.

## Materials and Methods

### Animal studies

All breeding, care, husbandry and procedures were approved by The University of Pittsburgh Department of Laboratory and Animal Resources (DLAR) and the Institutional Animal Care and Use Committee (IACUC) with standard animal chow and water provided *ad libitum*. C57BL6 mice expressing GFP (C57BL/6-Tg(UBC-GFP)30Scha, MGI:3057178) have been previously described and were used as a source of wild-type (WT) control hepatocytes due to the ease with which the GFP gene could be identified [25]. C57BL6 *c*-*myc*^*LoxP/LoxP*^ mice (B6.129S6-Myc^tm2Fwa^, MGI:2178233) have been previously described (Fig. 1A) [25, 37, 91, 92] and were obtained as a gift from I. Moreno de Alboran. The generation of mice bearing a 1717 bp deletion spanning exons 3-6 of the *Mlx* locus (*Mlx*KO mice) (Fig. 1B) has also been recently described [8]. Transgenic mice expressing a fusion protein comprised of the hormone-binding domain of the estrogen receptor and Cre recombinase (CreER) and under the control of the albumin promoter that allows CreER to be activated in hepatocytes following tamoxifen exposure (B6.129S2-Alb^tm1(cre/ERT2)Mtz^, MGI:3052812) were a kind gift of Dr. Frank Gonzalez (Laboratory of Metabolism, Center for Cancer Research National Cancer Institute). The latter mice were bred to homozygosity with *Mlx*^LoxP/LoxP^ mice or *Myc*^*LoxP/LoxP*^x *Mlx*^*LoxP/LoxP*^ mice [25, 37]. At weaning, mice were subjected to five daily i.p. injections of tamoxifen (75 mg/Kg each) in corn oil (Sigma-Aldrich, St. Louis, MO). Several weeks later, hepatocytes were harvested as previously described [25, 37]. An aliquot of these was used for DNA isolation and to quantify the extent of *Myc* and/or *Mlx* knockout (Fig. 1). The remainder of the *Myc*KO, *Chrebp*KO or *Myc*KO x *Chrebp*KO hepatocytes were then combined in indicated proportions with WT hepatocytes and a total of 3×10^5^ cells were injected intrasplenically into *Fah-/-* FRG-NOD mice (Yecuris, Inc., Tualatin, OR)[25, 37] (*ChreBP* mice: B6.129S6-Mlxipl^tm1Ku^, MGI:3043871; *Fah* mice: NOD.Cg-*Rag1*^*tm1Mom*^ *Fah*^*em1Mvw*^ *Il2rg*^*tm1Wjl*^ MGI:5485380). All animals were maintained on 8 mg/L NTBC (Ark Pharm, Libertyville, IL) in their drinking water. After four days, NTBC was discontinued until mice lost approx. 20% of their body weight. NTBC was then re-instated until mice regained their age-appropriate weight. NTBC cycling was continued either until mice had become NTBC-independent (at least 20 wk post-transplantation) or until wk 28 in those cases where NTBC-independence was not achieved. Hepatocyte DNAs were then isolated from recipients and the TaqMan based approaches shown in Table 1 and Fig. 1 were again used to determine the donor:recipient ratio and the relative contribution of each donor population [25, 37]. PCRs were performed in a volume of 12 μl with 50 ng of genomic DNA. Conditions for amplification were 95 °C for 5 min, 40 cycles at 95 °C for 15 s, 60 °C for 60 s.

For gene expression profiling, the above-described *Myc*^*LoxP/LoxP*^, *Mlx*^*LoxP/LoP*^ and *Myc*^LoxP/LoxP^x *Mlx*^*LoxP/LoP*^ mice were bred to B6.129-*Gt(ROSA)26Sor*^*tm1(cre/ERT2)Tyj*^/J mice (MGI:3699244), which expresses CreER under the control of the ubiquitously-expressed ROSA26 promoter (Jackson Labs) [93]. Excisional inactivation of each locus was initiated at the time of weaning and confirmed as described above. Liver RNAs were then obtained from mice that were the same age as those used for hepatocyte transplants (∼5 mos).

### Triglyceride assays

Total lipid was extracted from ∼50 mg of liver using the Folch method [94]. Total triglyceride content was then determined as described previously using the Free Triglyceride Reagent (Sigma-Aldrich, Inc.) [25, 95].

### Histology, immuno-histochemistry and immuno-blotting

Fresh tissues sections were immediately fixed in formalin, embedded in paraffin and stained with hematoxylin-eosin (H&E) as previously described [25, 37, 49]. Oil Red O (ORO) staining and immuno-histochemistry on snap-frozen sections were also performed as previously described [25, 37]. Tissue samples for immuno-blotting were disrupted in SDS-PAGE lysis buffer containing protease and phosphatase inhibitors but lacking β-mercaptoethanol or Bromophenol Blue as previously described [25, 46, 49, 71]. Protein quantification was performed using the BCA reagent according to the directions of the supplier (Sigma-Aldrich). Following β-mercaptoethanol (1%) and Bromophenol Blue (10%) addition, samples were boiled for 5 min, dispensed into small aliquots and stored at -80C until ready for use. SDS-PAGE and semi-dry transfer to PVDF membranes (Sigma-Aldrich) was performed as previously described [25, 46, 49, 71]. Antibodies used for immuno-blotting included rabbit monoclonals directed against Mlx and Myc (#85570 and #13987, Cell Signaling Technologies, Inc., Danvers, MA) and a mouse monoclonal antibody against GAPDH (#G8795, Sigma-Aldrich). A mouse monoclonal anti-Ki-67 antibody used for immuno-histochemistry was also from Cell Signaling Technologies (#12202). Horseradish peroxidase secondary antibodies were from Santa Cruz Biotechnology (Santa Cruz, CA). Ki-67 immunostain quantification was performed using the ImageJ Immunohistochemistry (IHC) Image Analysis Toolbox (https://imagej.nih.gov/ij/plugins/ihc-toolbox/index.html). All antibodies were used at the dilutions recommended by the suppliers. Immunoblots were developed using an enhanced chemiluminescent assay kit as directed by the supplier (SuperSignal™ West Pico Plus, Thermo-Fisher, Inc., Waltham, MA).

### RNAseq and ChIP-seq experiments

RNAs were purified from five replica tissues of each group of mice using Qiagen RNeasy Mini Kit (Qiagen, Inc. Germantown, MD) followed by DNase digestion [25, 37]. RNA concentration and integrity was confirmed on an Agilent 2100 Bioanalyzer (Agilent Technologies, Foster City, CA) and only those samples with RIN values of >8.5 were used for sequencing. All subsequent analyses were performed as previously described [25, 71]. Sample preparation and sequencing was performed on a NovaSeq 600 instrument (Illumina, Inc., San Diego, CA) by Novagene, Inc. (Sacramento, CA) and raw data were deposited in the National Center for Biotechnology Information (NCBI) Gene Expression Omnibus data base (GEO) (Accession no GSE181371). Data sets from previous RNAseq studies of *Myc*KO, *Chrebp*KO and *Myc*KO x *Chrebp*KO livers and mutant forms of β-catenin+YAP^S127A^ HBs are available from the GEO data base sets GSE114634 and GSE130178 [25, 71]. Differential gene expression was assessed by three different approaches, namely EdgeR, CLC Genomics Workbench version 21 (Qiagen) and DeSeq2, as previously described [25, 47]. When low-abundance reads (cpm <1) were encountered for both comparisons, they were eliminated. Reads from FASTQ files were mapped to the GRCm38.p6 mouse reference genome using STAR (https://github.com/alexdobin/STAR/releases) version 2.7.5. BAM-formatted output was analyzed and transcript abundance was determined by featureCounts (http://bioinf.wehi.edu.au/featureCounts/). Where necessary, Ingenuity Profiling Analysis (IPA) (Qiagen) was used to classify transcripts into pathways whose significance was adjusted for false discovery using the Bonferonni–Hochberg correction. We further utilized Gene Set Enrichment Analysis (GSEA) [48] to identify alterations of functionally-related groups of transcripts from the Molecular Signatures Database (mSigDB) C2 collection (v.7.4) (http://www.gsea-msigdb.org/gsea/msigdb/index.jsp) or from the MitoProteome Data base (http://www.mitoproteome.org). Volcano plots were generated using the R software package ggplot2 (https://ggplot2.tidyverse.org/) with significant differences between samples being defined as having fold differences >1.5 and false discovery rates <0.05. Heat maps were generated using the ComplexHeatmap package (version 2.6.2, https://bioconductor.org/packages/release/bioc/html/ComplexHeatmap.html). Statistical analyses were performed with R software v4.0.3 (R Foundation for Statistical Computing, Vienna, Austria) and GraphPad Prism v9.00 (GraphPad Software Inc., San Diego, CA).

To analyze ChIP-seq data, we explored the binding Myc and Mlx to their target gene sequences in two different HepG2 cell lines that had been modified using Crispr so as to introduce 3 x FLAG tags at the C-termini of each protein. This allowed ChIP-seq to be performed under identical conditions with a single anti-FLAG antibody. The results were down-loaded from the ENCODE website (https://www.encodeproject.org/) and analyzed using ChIPpeakAnno version 3.13 and the annotation datbase TxDb.Hsapiens. UCSC.hg38.knownGene (R package version 3.13.0.)[96]. Only binding sites residing within +/-2.5 kb of the transcriptional start of each target gene were considered for the current analysis[53, 97]. Overlap between Myc and Mlx binding regions was obtained using the **“**findOverlapsOfPeaks” funcation (set maxgap = 0, minoverlap = 0). Venn diagrams were used to display unique and overlapping binding sites. FIMO (Version 5.4.1) from the MEME software suite was used to identify E-boxes and ChoREs most closely associated with ChIP-seq peaks. Categorization of genes associated with bound peaks was performed using previous described collections of functionally-related genes or those from the IPA and mitoproteome data bases (Qiagen, Inc. and http://www.mitoproteome.org).

## Acknowledgements

Analysis of RNAseq data was supported by The University of Pittsburgh Center for Research Computing. This work was supported by NIH grant RO1 CA174713 and by a Hyundai Hope on Wheels Scholar grant to EVP.

